# Hyperlipidemia drives tumor growth in a mouse model of obesity-accelerated breast cancer growth

**DOI:** 10.1101/2025.02.10.637542

**Authors:** Renan FL Vieira, Sawyer R Sanchez, Menusha Arumugam, Peyton D Mower, Meghan C Curtin, Molly R Gallop, Jillian Wright, Alexis Bowles, Gregory S Ducker, Keren I Hilgendorf, Amandine Chaix

## Abstract

Obesity is an established risk factor for breast cancer (BC), yet the specific mechanisms driving this association remain unclear. Dysregulated lipid metabolism has emerged as a key factor in cancer cell biology. While obesity is often accompanied by hyperlipidemia, the isolated impact of elevated lipid levels on BC growth has not been experimentally tested. Using the E0771 orthotopic model of obesity-accelerated BC growth in immune-competent mice, we investigated the direct role of systemic lipids in tumor growth. Combining dietary and genetic mouse models, we show that elevated circulating lipids are sufficient to accelerate BC tumor growth even in the absence of obesity or alterations in blood glucose and/or insulin levels. Pharmacological lowering of systemic lipid levels attenuates BC growth in obese mice, suggesting a direct role for lipids in fueling tumor expansion. Notably, we also show that weight loss alone, without a corresponding reduction in lipid levels such as that induced by a ketogenic diet, fails to protect against BC, highlighting the necessity of targeting lipid metabolism in obesity-associated BC. Our findings establish hyperlipidemia as a critical driver of BC progression and suggest that lipid-lowering interventions may be a promising strategy to mitigate BC risk in obese individuals.

## Introduction

Obesity is associated with an increased incidence of 13 cancers, including breast cancer (**BC**) in postmenopausal women.^1,2^ Breast cancer affects 1 in 8 women in the USA and each 5 kg/m^2^ increase in body mass index (**BMI**) is associated with a 12% increase in breast cancer risk.^3,4^ Obesity is a complex condition associated with changes to both the systemic and local microenvironment of the breast tumor. As such, multiple mechanisms likely act in concert to promote tumor growth which complicates the identification of specific cancer drivers. Increased adiposity is often accompanied by adipose tissue dysfunction that may directly influence breast tumors growing in the mammary fat pad. Obesity is also associated with dysregulated circulating hormones and metabolites including insulin, glucose, and lipids, often culminating in metabolic syndrome.^5–7^ Consequently, tumors that arise in patients with obesity are exposed to a fundamentally different nutrient environment than those from lean patients and mounting evidence suggests they adopt a different metabolic program.^8–10^ With half of adults globally, and 70% of adults in the United States having a BMI >25^11^, it is critical to identify specific mechanisms that lead to accelerated tumor growth in individuals who are overweight or have obesity and thus tailor therapeutic and preventive approaches around a patient’s metabolic status.

Obesity-accelerated breast tumor growth can be modeled in female C57BL/6 mice using the syngeneic triple-negative BC cell line E0771 injected orthotopically into the mammary fat pad. In this model, tumor growth is significantly accelerated upon injection into obese mice fed a 60% high fat diet (**HFD**) compared to low fat diet (**LFD**)-fed lean mice.^12–15^. Using this model, studies have shown that different discrete metabolic alterations associated with obesity are sufficient to promote E0771 growth. Specifically, lowering systemic glucose levels by using SGLT2 inhibitors can reduce E0771 growth in an insulin-dependent manner in hyperglycemic animals^15^ and increased insulin was sufficient to enhance E0771 growth.^12^ Separately, excess adiposity is also sufficient to accelerate tumor growth^14^ with adipocyte-specific secretion of polar metabolites playing a critical role in driving tumorigenesis.^13,16^ However, the role of systemic hyperlipidemia, a key component of the pathophysiology of obesity, has not been investigated in this model system.

Lipids are essential sources of energy and building blocks that support tumor growth.^17^ In particular, tumors require a significant flux of energy to synthesize the fatty acid precursors of cellular membrane phospholipids to support uncontrolled cell division.^18,19^ This metabolic constraint suggests that tumors could streamline their metabolic program by increasing their uptake of exogenous lipids from the diet and adipose tissue when available.

In this study, we leveraged the power of the C57BL/6 mouse model’s genetic and metabolic response to dietary fat to evaluate the contribution of hyperlipidemia to obesity-accelerated BC growth. Using both diet and genetics to induce hyperlipidemia in the absence of hyperglycemia and hyperinsulinemia, we show that elevated serum lipid levels are sufficient to accelerate E0771 BC tumor growth irrespective of adiposity. We further show that hyperlipidemia is required for accelerated E0771 tumor growth in HFD-fed mice. Accordingly, weight loss strategies that do not suppress hyperlipidemia fail to retard E0771 tumor growth.

## Results

### 1 Several metabolic alterations are associated with the accelerated growth of E0771 BC cells in diet-induced obese female mice

To interrogate the role of dietary lipids in accelerating BC tumor growth, we used the well-characterized E0771 syngeneic BC tumor model. 6–8-week-old female C57BL/6J mice were fed a lard-based HFD with 60% kcal from fat (Research Diet, D12492) or a LFD with 10% kcal from fat. Body weight (**BW**) gain was significantly higher in HFD-compared to LFD-fed females, with the average body weight diverging as early as 2 weeks upon study start (Fig.1.A, average of 2 independent experiments). Weekly monitoring of body composition in a subset of mice during the first 4 weeks revealed a sharp increase in the percentage of fat mass following one week of HFD feeding, followed by a steady progressive increase during the following weeks (Fig.1.B). In contrast, the percentage of fat mass did not change over time in LFD-fed mice (Fig.1.B). After 7-8 weeks, females on HFD had on average twice the body fat mass percentage as mice on LFD (Fig.1.C, 2 independent experiments).

**Figure 1:**
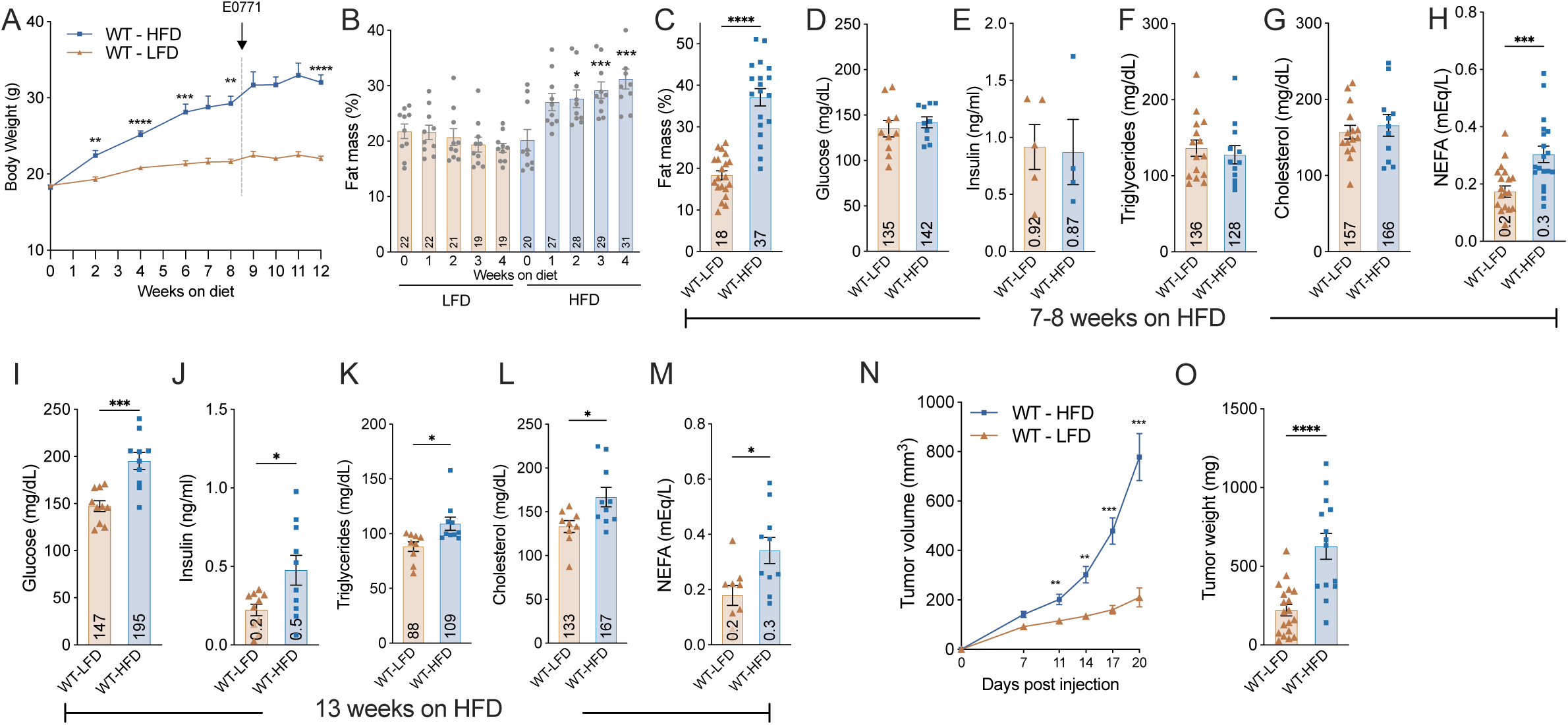
E0771 BC tumor growth is accelerated in female mice fed a 60% HFD. **A**. Body weight over time in WT female C57BL/6J mice (6-9 weeks old at start) fed a LFD (brown triangles, n=20, pool of 2 independent cohorts) or HFD (blue squares, n=17, pool of 2 independent cohorts). E0771 are injected orthotopically in both the left and the right 4^th^ mammary fat pad after 8-9 weeks on diets. **LFD:** low fat diet, 10% kcal from fat). **HFD:** high fat diet, 60% kcal from fat. **B, C**. Total fat mass as (B) the percentage of total body weight during the first 4 weeks on diet and (C) average % fat mass of 3 independent cohorts after 7 to 8 weeks on diet prior to E0771 injections. **D-H**. Blood (D) glucose, serum (E) insulin, (F) triglycerides, (G) cholesterol and (H) NEFA after 7-8 weeks on diet. **I-M**. Fasting (I) blood glucose, serum (J) insulin, (K) triglycerides, (L) cholesterol and (M) fed NEFA after 13 weeks on diet. **N, O**. Tumor volume over time (N) and weight at experimental endpoint (day 21) (O). <u>Statistics</u>: A. REML mixed-effects model with Sidak’s multiple comparisons post hoc test. Significance is shown as * for LFD vs HFD. B,N. Two-way repeated measures ANOVA with Sidak’s multiple comparisons post hoc test. C-M, O. Unpaired t test. Individual data points are represented together with mean and sem. For all, * p<0.05, ** p<0.01, *** p<0.001.

Systemic metabolic alterations such as increased serum glucose, insulin, and lipids are well-established in C57BL/6J males fed a 60% HFD. Females however are known to be more resistant.^20–22^ Accordingly, we did not observe significant changes in glucose, insulin, triglycerides (**TG**), or cholesterol (**Cho**) between LFD- and HFD-fed females after 7-8 weeks of feeding (Fig.1.D-G). Fed serum levels of non-esterified fatty acids (**NEFA**) were significantly elevated in HFD- versus LFD-fed mice as early as 7-8 weeks (Fig.1.H). Markers of metabolic dysfunction were reassessed in after 13 weeks on HFD. At this time, we observed elevated serum levels of glucose, insulin, TG and Cho in HFD-fed mice specifically in the fasted state (Fig.1.I-L). NEFA were still elevated in the fed state (Fig.1.M). These results suggest that metabolic dysfunction develops over time in female C57BL/6J, albeit to a slower and lesser extent than commonly reported in their male counterparts.

Confident that HFD-fed female mice were developing classic hallmarks of metabolic dysfunction between 8 and 13 weeks on diet, we initiated tumor studies in animals after 8-9 weeks on HFD. The triple negative BC cell line E0771 was injected in the 4th mammary fat pad of the mice and tumors monitored for development over 3 weeks. Tumor growth was significantly accelerated (Fig.1.N) in HFD-fed mice and final tumors weights were increased 3-fold (626 mg in HFD vs 220 mg in LFD) after 3 weeks (Fig.1.O).

### 2 Elevated serum lipids are sufficient to accelerate E0771 growth in mouse models of genetic hyperlipidemia

To be able to determine the contribution of hyperlipidemia to the accelerated growth of E0771 tumors in the absence of excess adiposity, we leveraged two genetic models of hyperlipidemia, the apolipoprotein E knock out mouse (**ApoE KO** JAX #002052)^23^ and the low density lipoprotein receptor knock out mouse (**LDLR KO**, JAX #002207)^24^. In a first experiment, mice were fed a western diet (**WD**) with 40% kcal from fat and 1.25% cholesterol (D12108C) which is the diet classically used in atherosclerosis studies and known to rapidly induce hypertriglyceridemia and hypercholesterolemia in ApoE KO and LDLR KO mice.^25–29^ 6 weeks of WD feeding in WT females led to a modest increase in BW associated with a small increase in fat mass when compared to LFD feeding (Fig.2.A, 2.B, 24% vs 32% fat after 6 weeks). ApoE KO and LDLR KO mice on WD did not gain weight and were significantly leaner than WT controls (Fig.2.A, 2.B, 17% and 18% body fat for ApoE KO and LDLR KO respectively). Mice in all groups had normal levels of insulin and glucose (Fig.2.C, 2.D). As expected, ApoE KO and LDLR KO mice were hyperlipidemic. Serum TG and Cho levels were between 2-10 fold higher in both ApoE KO and LDLR KO females compared to WT females fed LFD or WD (Fig.2.E, 2.F). Serum NEFA levels were not different between groups (Fig.2.G). E0771 cells were injected bilaterally in the 4^th^ mammary breast fat pads of the mice after 4 weeks on their respective diets and tumors allowed to grow for 3 weeks. E0771 tumors were significantly larger in ApoE KO and LDLR KO compared to WT females fed either LFD and WD (Fig.2.H). Compared to WT females on WD, tumors in LDLR KO mice were twice as large and tumors in ApoE KO almost three times larger (Fig.2.H).

**Figure 2:**
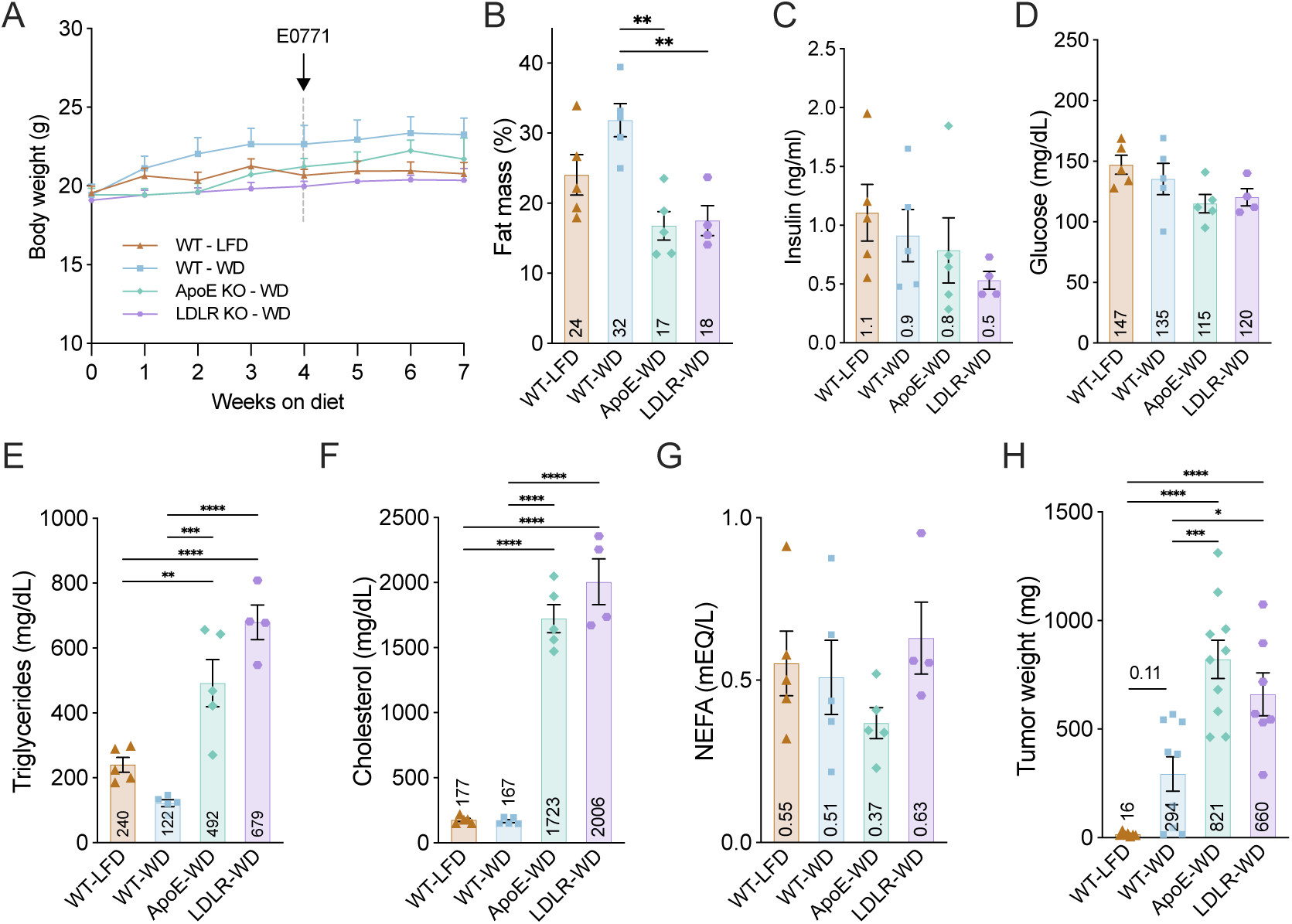
E0771 tumor growth is accelerated in ApoE KO and LDLR KO mice fed a western diet. **A.** Body weight of female WT C57BL/6J mice fed a WD (blue squares, n=5), ApoE KO on WD (green diamonds, n=5), and LDLR KO on WD (purple circles, n=4), and WT females fed a LFD (brown triangles, n=5). 8-week-old mice were placed on diets for 4 weeks before E0771 cells injection and maintained on their diet during the 3 weeks of breast tumors growth. **LFD:** low fat diet, 10% kcal from fat. **WD:** western diet, 40% kcal from fat with 1.25% cholesterol. **B**. Fat mass as the percentage of total body weight after 6 weeks on diets. **C-G**. Serum level of (C) insulin, (D) blood glucose measured with a glucometer, serum (E) triglycerides, (F) cholesterol and (G) NEFA all in the randomly fed state 5 days before E0771 injections. **H**. Tumor weights 3 weeks post injections. Data include the weights of all bilateral tumors that did not clear. (n=6 for WT-LFD, n=9 for WT-WD, n=10 for ApoE-WD and n=7 for LDLR-WD) <u>Statistics</u>: A. Two-way Repeated measures ANOVA with Tukey’s multiple comparisons post hoc test. B-H. One-way ANOVA with Tukey’s multiple comparisons post hoc test. Individual data points are represented together with mean and sem. For all, * p<0.05, ** p<0.01, *** p<0.001.

In a second experiment to directly compare the effects of hyperlipidemia on tumor growth in the context of obesity, we initiated E0771 tumors in ApoE KO versus WT females fed a 60% HFD. Consistent with prior cohorts, 60% HFD feeding led to a significant increase in BW as early as 1 week on the diet and a significant increase in fat mass compared to LFD feeding in WT females (Fig.3.A, 3.B). ApoE KO mice on HFD weighed more than LFD fed WT mice after 6 weeks on the diet but they remained significantly lighter than WT females (Fig.3.A) but with a similar percentage of fat mass as WT mice on LFD (Fig.3.B). Mice in all groups had similar levels of insulin, glucose and NEFA after 6-7 weeks on their respective diets (Fig.3.C, 3.D, 3.G). In contrast, TG levels were twice as high and Cho levels three times higher in ApoE KO females compared to WT females on either diet (Fig.3.E, 3.F). Like what we observed with a WD, E0771 tumor growth was accelerated in ApoE KO on a HFD compared to WT on HFD (Fig.3.H). These data show that hyperlipidemia alone, without obesity nor alterations in glucose or insulin, is sufficient to accelerate the orthotopic growth of E0771 tumors.

**Figure 3:**
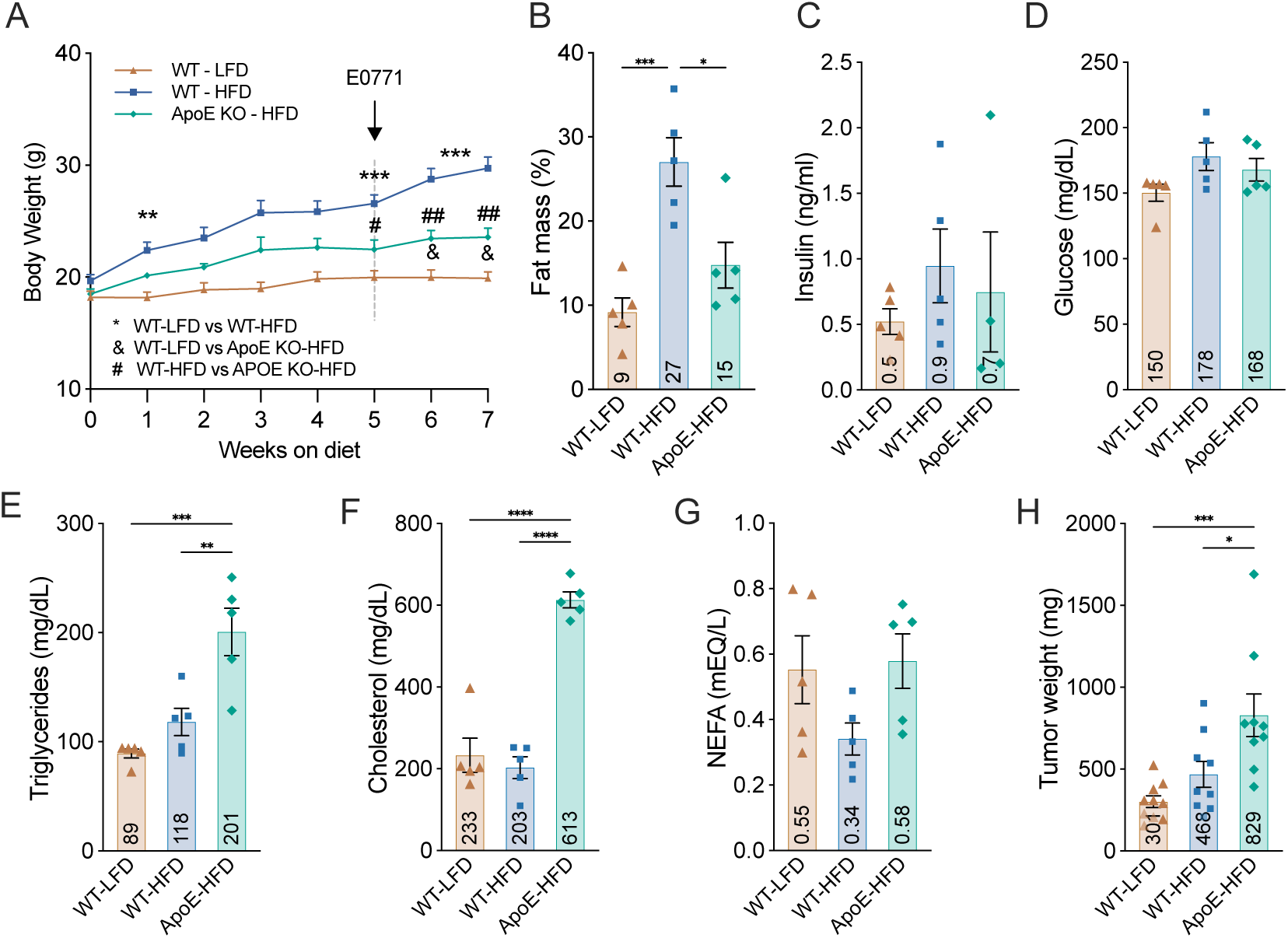
E0771 growth in 60% HFD-fed WT and ApoE KO mice. **A.** Body weight of female WT C57BL/6J mice on HFD (blue squares, n=5), ApoE KO on HFD (green diamonds, n=5) and WT on LFD (brown triangles, n=5). 7-week-old mice were placed on diets for 5 weeks before E0771 cells injection and maintained on their diet during 3 weeks of tumor growth. **LFD:** low fat diet, 10% kcal from fat). **HFD:** high fat diet, 60% kcal from fat. **B**. Fat mass as the percentage of total body weight after 4 weeks on diets. **C**. Serum fasting level of insulin prior to E0771 injections. **D-G**. Blood glucose (D), serum (E) triglycerides, (F) cholesterol and (G) NEFA in the randomly fed state prior to E0771 injections. **H**. Tumor weights at endpoint 3 weeks post injections. Data include the weights of all bilateral tumors that did not clear. (n=10 for WT-LFD, n=9 for WT-HFD, n=9 for ApoE-WD) <u>Statistics</u>: A. Two-way repeated measures ANOVA with Tukey’s multiple comparisons post hoc test. Significances are shown as * for WT-LFD vs WT-HFD, & for WT-LFD vs ApoE KO-HFD and # for WT-HFD vs APOE KO-HFD. B-H. One-way ANOVA with Tukey’s multiple comparisons post hoc test. Individual data points are represented together with mean and sem. For all, * p<0.05, ** p<0.01, *** p<0.001.

### 3 Diet-induced hyperlipidemia is sufficient to accelerate E0771 growth

In prior work, we observed that C57BL/6J female mice fed a 90% fat ketogenic diet (**KD**) for extended periods of time will develop obesity and systemic hyperlipidemia, but are protected from the hyperglycemia and hyperinsulinemia observed in adiposity-matched 60% HFD-fed females.^22^ We asked whether this diet-mediated hyperlipidemia was also sufficient to accelerate the growth of E0771 tumors. 8-week-old female C57BL/6J mice were fed a 90% KD or a control LFD for 10 weeks. BW gain in KD-fed mice was associated with a progressive increase in adiposity (Fig.4A, 4B). Insulin levels were not different between LFD- and KD-fed females and blood glucose were unchanged in fasted mice and significantly lower 4 hours after refeeding in KD mice compared to LFD-fed mice. (Fig.4C, 4D). Conversely, postprandial TG levels were 3.5 times higher, Cho levels 2.5 times higher and NEFA two times higher in KD-fed mice than LFD-fed mice (Fig.4E, 4F, 4G). E0771 cells were injected in the breast fad pads after 10 weeks of dietary intervention. After 3 weeks of cell growth, E0771 tumors were twice as large in KD mice (Fig.4H, average of 330 mg in LFD- versus 695 mg in KD-fed mice).

**Figure 4:**
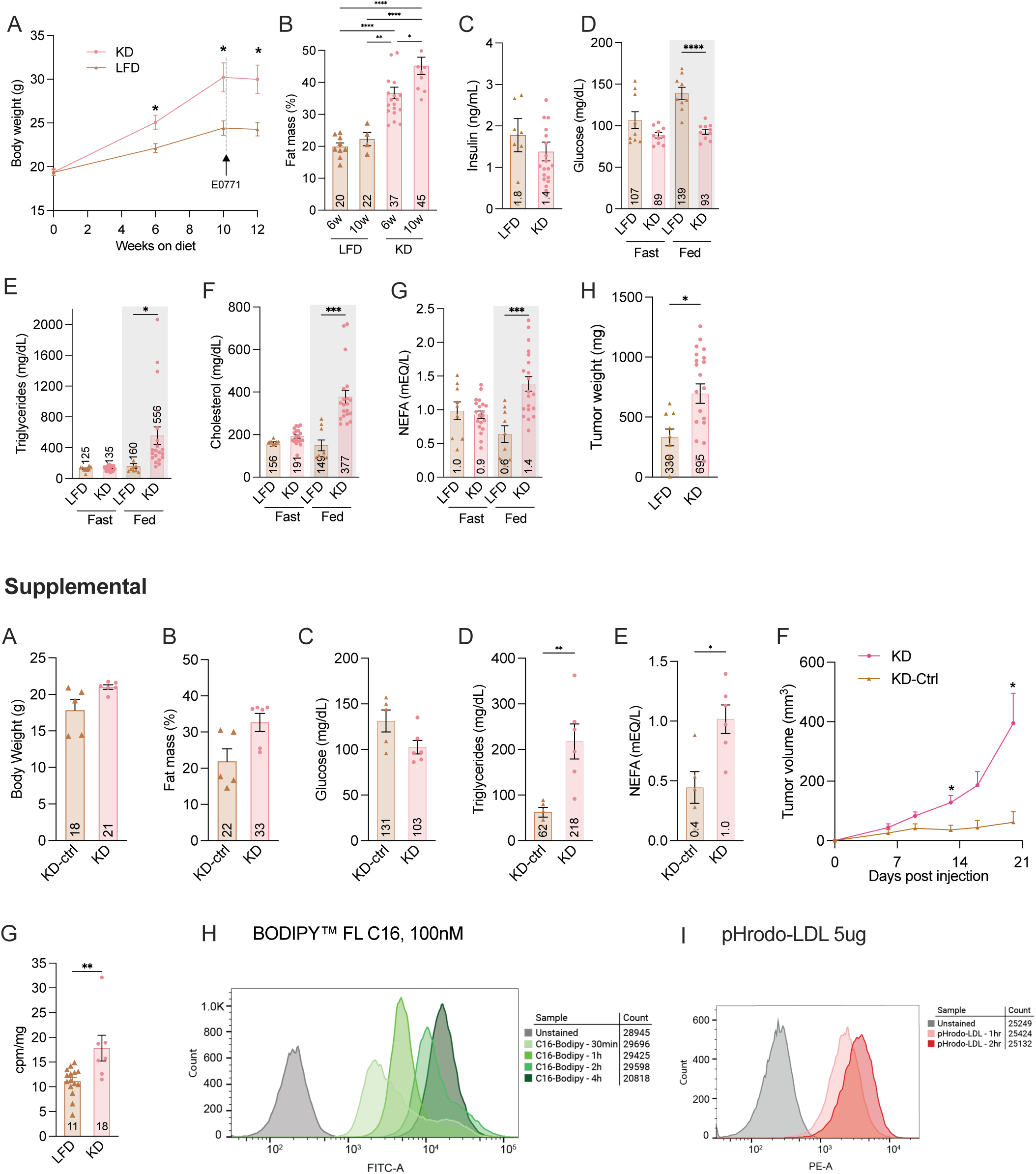
E0771 tumor growth can be accelerated by 90% fat ketogenic diet. **A.** Body weight of female WT C57BL/6J mice fed a 90% fat KD (pink circles, n=20) or a LFD (brown triangles, n=9). 8-week-old mice were placed on diets for 10 weeks before E0771 cell injection and maintained on their diet during the 3 weeks of tumor growth. **LFD:** low fat diet, 10% kcal from fat. **KD:** ketogenic diet, 90% kcal from fat. **B**. Fat mass as the percentage of total body weight after 6 and 10 weeks on diets. **C**. Serum fasting level of insulin prior to E0771 injections. **D-G**. Fasted and 4h refed blood glucose measured with a glucometer (D) and serum levels of (E) triglycerides, (F) cholesterol, and (G) NEFA. **H**. Tumor weights at endpoint 3 weeks post injections. Data include the weights of all bilateral tumors that did not clear. (n=9 for WT-LFD, n=21 for WT-KD) <u>Statistics</u>: A. REML mixed-effects model with Sidak’s multiple comparisons post hoc test. Significance is shown as * for LFD vs KD. B. One-way ANOVA with Tukey’s multiple comparisons post hoc test. C, H. Unpaired t test. D-G. Unpaired t test, within fast or fed condition. Individual data points are represented together with mean and sem. For all, * p<0.05, ** p<0.01, *** p<0.001. <u>Supplemental</u>: **S4A**. Body weight of WT females fed a 90% fat KD (pink circles, n=6) or a low fat KD-Ctrl (brown triangles, n=5) before E0771 injections. 8-week-old mice were placed on diets for 4 weeks before E0771 cells injection and maintained on their diet during the 3 weeks of breast tumors growth. **KD-ctrl:** low fat diet, 10% kcal from fat and 10% kcal from proteins. **KD:** ketogenic diet, 90% kcal from fat. **S4B**. Fat mass as the percentage of total body weight after 4 weeks on diets. **S4C-E**. Blood glucose (C), serum (D) triglycerides, and (E) NEFA in the randomly fed state prior to E0771 injections. **S4F**. Tumor volume over time. **S4G**. Total ^3^H radioactive counts in tumors 16 h post ^3^H-triolein bolus. **S4H, I**. Fluorescence intensity in E0771 cells treated with (H) 100nM of C16-Bodipy and (I) 1uM of LDL-pHrodo™ Red. <u>Statistics</u>: A-E,G. Unpaired t test. F. Two-way repeated measures ANOVA with Sidak’s multiple comparisons post hoc test. Significance is shown as * for KD-ctrl vs KD.

To control for protein content between conventional LFD and KD, we repeated our study using a LFD with 10% kcal from protein rather than 20% to match the protein content of the KD used in these studies (KD-ctrl diet). Additionally, we injected the E0771 cells before a significant divergence in BW and % body fat had occurred between control and KD-fed mice (Fig.S4A, S4B). 4 weeks of KD feeding was sufficient to induce significant increases in serum TG and Cho without differences in glucose (Fig.S4C-E). The growth of E0771 breast tumors was greatly accelerated by a KD in this experiment (Fig.S4F, 62 mg vs 395 mg). To confirm that E0771 cancer cells in hyperlipidemic mice take up more lipids than in LFD-fed mice, we administered a bolus of ^3^H-triolein (([9,10-^3^H(N)]-triolein) by oral gavage. Radioactive counts were measured 16 hours later in KD and LFD tumors. E0771 cells in KD mice took up significantly more ^3^H signal than controls (Fig.S4G). We independently confirmed the ability of E0771 cells to readily take up both NEFA and lipoproteins *in vitro* (Fig.S4H, S4I). These results suggest that diet-induced hyperlipidemia, without hyperinsulinemia or hyperglycemia, is sufficient to accelerate BC growth.

### 4 Systemic lipid lowering prevents the obesity-accelerated growth of E0771 tumors

Having established that systemic hyperlipidemia can accelerate the growth of E0771 tumors injected into the mammary fat pad, we set out to test whether lowering blood lipids levels would prevent the obesity accelerated growth of E0771 cells. To lower blood lipid levels, we used an antisense oligonucleotide (ASO) targeting angiopoietin-like 3 (Angptl3; Ionis). Treatment with this ASO reduces hepatic expression of Angptl3 resulting in lower TG and Cho in C57BL/6J mice.^30^ Female mice were fed a control LFD or a 60% HFD for 10 weeks to induce obesity (Fig.5.A). Prior to the start of the treatment, fed blood was collected and serum TG and Cho levels measured to randomize mice within each diet into 2 groups of equivalent TG and Cho (Fig.5.B, 5.E). One group received the ASO targeting Angptl3 (Angpt3-ASO) while the control group received a scramble ASO (Scr-ASO). Mice were injected subcutaneously with 25 mg/kg on days 1, 3 and 7 of the first week and then weekly for a total of 9 injections (Fig.5.A). While there was a small and non-significant difference in BW between the two HFD groups at the start of the treatment, neither ASO injection affected weight gain trajectories and HFD-fed mice continued to gain weight throughout the study (Fig.5.A). At terminal collection, adiposity was similar between Angpt3-ASO-treated and Scr-ASO-treated mice (Fig.S5A). Serum analysis of TG and Cho showed significant reductions in TG and Cho in both the LFD and HFD groups treated with Angpt3-ASO versus scramble after 4 injections (2 weeks after the start of the treatment) (Fig.5.C, 5.F). NEFA levels were not different (Fig.S5B). Reduced levels of TG and Cho were maintained throughout the experiment as measured at the terminal collection (Fig.5.D, 5.G), while glucose and insulin levels were not affected by the treatment (Fig.S5C, S5D). E0771 cells were injected 14 weeks after diet initiation and 5 ASO injections (Day 96). Tumor size measured by caliper revealed reduced E0771 tumor growth in HFD-fed mice treated with the Angptl3-ASO compared to scramble (Fig.5.H). At endpoint, tumors in the HFD group treated with the Angptl3-ASO were 1.6 times smaller than in the HFD control group that received the scramble ASO (1628 mg vs 1029 mg). Angptl3-ASO treatment had no effect on tumors in LFD-fed mice (Fig.5.I). These results suggest that in this model of obesity-accelerated BC growth, lowering blood lipids is sufficient to retard tumor growth even when adiposity is unchanged.

**Figure 5:**
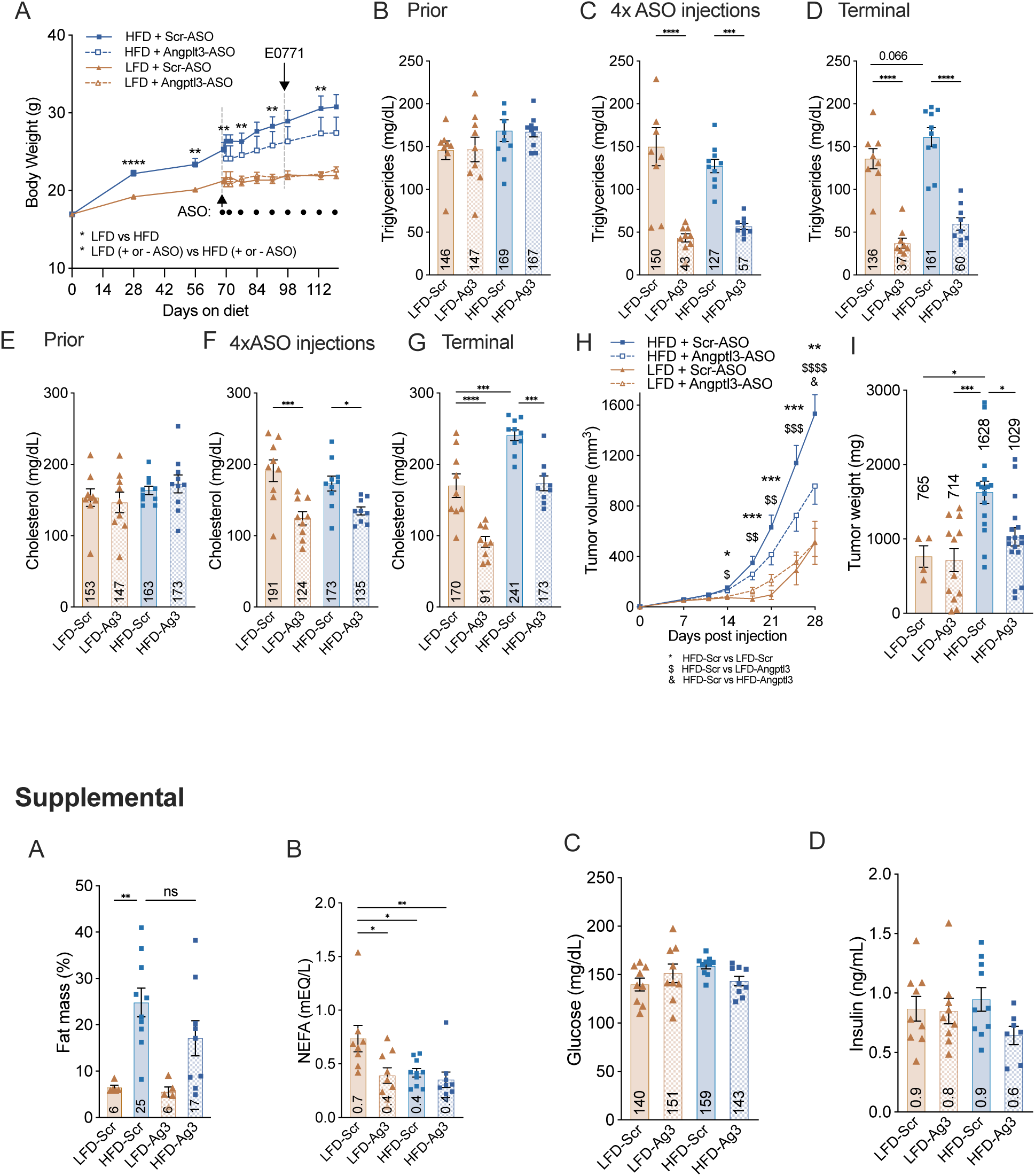
Systemic lipid lowering attenuates obesity-accelerated E0771 tumor growth. **A.** Body weight of female WT C57BL/6J mice throughout the experiment. **LFD:** low fat diet, 10% kcal from fat. **HFD:** high fat diet, 60% kcal from fat. 7-week-old mice were placed on their respective diets for 10 weeks before ASOs injections. Half the mice on HFD received the Angptl3-ASO (blue dotted line, triangles, n=10) and the other half the scramble-ASO (Scr-ASO, blue solid line, squares, n=10). Half the mice on LFD received the Angptl3-ASO (brown dotted line, diamonds, n=9) and the other half the scramble-ASO (brown solid line, triangles, n=9). Mice were treated for 4 weeks (5 ASO injections) before E0771 injections. Weekly treatment and experimental diets were maintained for the 3 weeks of breast tumor growth. ASOs were injected subcutaneously on day 1 (experimental day 70), day 3 and day 7 of the first week and then weekly at 25mg/kg. **B-G**. Serum levels of (B) triglycerides and (E) cholesterol prior to ASOs injections (experimental day 60), (C) triglycerides and (F) cholesterol after 4 ASOs injections (experimental day 85), and (D) triglycerides and (G) cholesterol at euthanasia (experimental day 125). **H, I**. Tumor volume over time (H) and (I) weight at experimental endpoint. <u>Statistics</u>: A. REML mixed-effects model with Tukey’s multiple comparisons post hoc test. Significances are shown as * for LFD vs HFD prior to ASO injections, and as * for LFD (both with Scr-ASO and Angptl3-ASO) vs HFD (both with Scr-ASO and Angptl3-ASO) since no significant differences were detected between Angptl3-ASO and Scr-ASO treatments within each diet group. B-G, I. One-way ANOVA with Tukey’s multiple comparisons post hoc test. H. REML mixed-effects model with Sidak’s multiple comparisons post hoc test. Significances are shown as * for HFD-Scr vs LFD-Scr, $ for HFD-Scr vs LFD-Angptl3 and & for HFD-Scr vs HFD-Angptl3. Individual data points are represented together with mean and sem. For all, * p<0.05, ** p<0.01, *** p<0.001. <u>Supplemental</u>: **A**. Fat mass as percentage of total body weight one day prior to collection (experimental day 124). **B,C,D**. Serum levels of (B) NEFA, (C) glucose and (D) insulin at euthanasia (experimental day 125). <u>Statistics</u>: A-D. One-way ANOVA with Tukey’s multiple comparisons post hoc test. Individual data points are represented together with mean and sem. For all, * p<0.05, ** p<0.01, *** p<0.001.

### 5 Weight loss without systemic lipid lowering does not reduce the growth of E0771 breast cancer tumors

The lipid dependency of our model of obesity-accelerated BC growth predicts that strategies to mitigate breast cancer recurrence need to focus on lipid-lowering beyond pure weight loss (**WL**). In our prior work, we showed that obese male mice switched to a 90% fat KD lost weight, and while this WL intervention reduced adiposity and normalized blood glucose and insulin, high lipid levels were maintained.^22^ To test how such a metabolic profile influences obesity-accelerated BC, we compared E0771 tumor growth between obese and previously obese females following two dietary interventions to induce WL: a KD and a LFD. Following 11 weeks of weight gain on HFD, mice were randomized to stay on HFD, or switched to either KD or LFD while control mice were maintained on LFD throughout (Fig.6A). Switching obese females to LFD led to a significant weight loss and normalization of fat mass (Fig.6A, 6B), while a KD prevented further weight gain and fat mass accrual (Fig.6A, 6B). E0771 cells were injected 6.5 weeks after initiation of the WL. At the time of injection, BW in all groups had stabilized (Fig.6A), and serum insulin levels and blood glucose were significantly lower in both WL groups compared to mice maintained on HFD, and not different than that of control mice maintained on LFD (Fig.6C, 6D). However, mice switched to KD maintained elevated TG, Cho and NEFA levels, with TG and NEFA being higher than even obese mice maintained on HFD (Fig.6E, 6F, 6G). Strikingly, while endpoint tumor mass was significantly lower in mice switched to LFD compared to HFD controls (Fig.6H), in mice who were switched to KD, E0771 tumor growth was not significantly different than HFD controls (Fig.6H) despite them being significantly leaner and having improved glucose homeostasis. Interestingly, although not statistically significant, the endpoint mass of E0771 tumors in the HFD->LFD group trended larger than in the always lean LFD group (Fig.6H), suggesting that metabolic alterations that develop obesity have long lasting consequences on BC, even when adiposity and serum metabolic profile is restored to that of lean mice. Overall, our weight loss data demonstrate again that adiposity per se is not the sole driver of tumor growth in obesity and that serum nutrient availability significantly influences tumor growth and metabolism.

**Figure 6.**
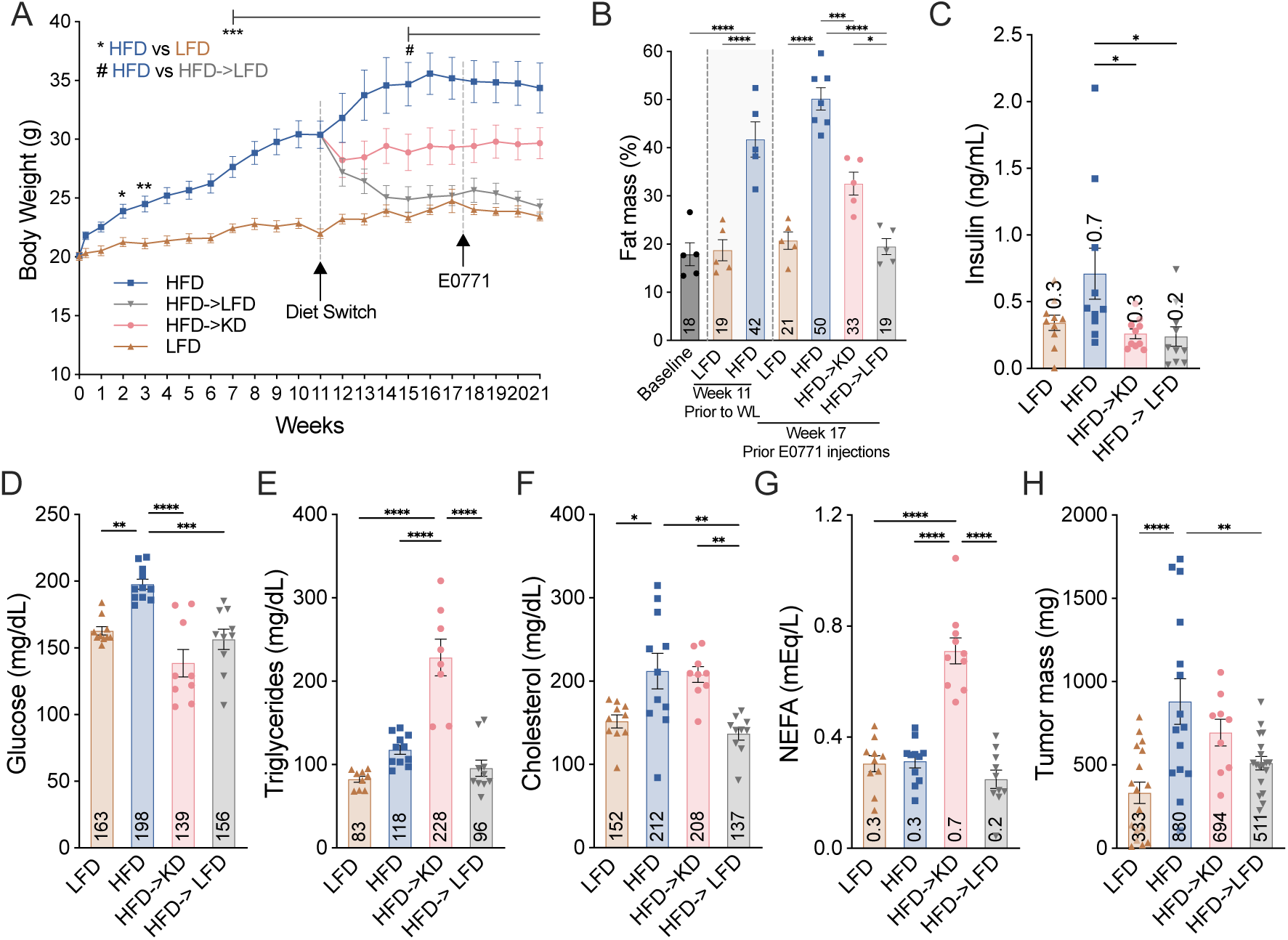
**A.** Body weight of female WT C57BL/6J mice throughout the experiment. 10-week-old females were placed on LFD or HFD for 11 weeks to induce weight gain prior to starting a weight loss (**WL**) intervention on KD. **LFD:** low fat diet, 10% kcal from fat. **HFD:** high fat diet, 60% kcal from fat. **KD**: ketogenic diet, 90% kcal from fat. Subsets of mice on HFD were either maintained on HFD (blue squares, n=10), or switched to a KD (HFD->KD, pink circles, n=9) or to a LFD (HFD->LFD, grey inverse triangles, n=10). Mice on LFD stayed on LFD (brown triangles, n=10). E0771 cells were injected 6.5 weeks after the start of the WL intervention and mice maintained on their dietary regimen during the 3.5 weeks of breast tumors growth. **B**. Fat mass as the percentage of total body weight for a subset of mice at baseline (experimental day 0), after 11 weeks of weight gain before the start of the WL intervention, and before E0771 injections (experimental week 17.5). **C**. Serum fasting insulin 2 days prior to E0771 injections **D-G**. Blood glucose (D), serum (E) triglycerides, (F) cholesterol and (G) NEFA at terminal collection. **H**. Tumor endpoint weights 24 days post injections. Data include the weights of all bilateral tumors that did not clear. (n=17 for LFD, n=15 for HFD, n=18 for HFD->LFD, n=9 for HFD->KD). <u>Statistics</u>: A. REML mixed-effects model with Sidak’s multiple comparisons post hoc test. Significances are shown as * for LFD vs HFD, and as # for HFD vs HFD->LFD. B-H. One-way ANOVA with Tukey’s multiple comparisons post hoc test. Individual data points are represented together with mean and sem. For all, * p<0.05, ** p<0.01, *** p<0.001.

## Discussion

### Hyperlipidemia is necessary and sufficient for accelerated breast cancer growth

In the US, more than 70% of adults are overweight or have obesity,^1,11,31^ a known risk factor for 13 cancers that accounts for 15-20% of all cancer-related deaths.^2^ Obesity, which affects 40% of US women, increases the incidence, morbidity, and mortality of postmenopausal breast cancer ^32^ and the risk of large, high-grade tumors, metastasis, and recurrence regardless of menopausal status.^3,4,33–40^ However, how obesity accelerates breast cancer remains unclear, in part because obesity causes many complex changes in the body such as increased adiposity and systemic metabolic alterations that can each directly influence BC growth.

Here, we show using both genetic and dietary approaches, that hyperlipidemia is sufficient to accelerate the growth of breast cancer in a mouse model, irrespective of adiposity and in the absence of hyperinsulinemia and hyperglycemia. This adds a significant new dimension to our understanding of how obesity can increase tumor growth. A major focus of cancer-obesity research has been centered on the role of loss of glycemic control in metabolic syndrome and how mechanisms to restore a normal glycemia can be antitumorigenic. These results have been most compelling in tumors with mutations in insulin signaling pathways (e.g. *PIK3CA*).^41^ In our tumor model without *PIK3CA* mutations, lowering systemic lipid levels significantly inhibited BC growth in obese mice without correcting the obesity. This suggests that BC cells in such an environment would likely be insensitive to de novo lipogenesis inhibitors and instead vulnerable to agents that target either systemic or local lipid availability. Targeting hyperlipidemia has therapeutic potential in breast cancer patients with obesity, representing approximately 50% of newly diagnosed cases.^42,43^ Finally, our data suggest that it is essential to consider hyperlipidemia in obesity-accelerated BC when orienting patients to weight loss strategies that might not all be equivalent in their ability to normalize systemic metabolic parameters and suppress BC recurrence.

### Lipids fuel tumor metabolism

Lipids are critical sources of biomass, energy, and inter- and intracellular signaling molecules for cancer cells, and changes in lipid abundance are a hallmark of many human tumors.^18,44^ Tumor-specific alterations in fatty acid synthesis, uptake, and oxidation help provide substrates for membrane construction and energy^45^, while changes in fatty acid saturation affect membrane fluidity as well as decrease tumor susceptibility to ferroptosis.^46^ Our data show increased lipid uptake by BC cells in hyperlipidemic conditions, but we do not know the fate of these lipids *in vivo*. *In vitro*, E0771 breast cancer cells readily take up both free fatty acids and lipoproteins. This points to potentially multiple mechanisms of lipid acquisition, or the ability to promiscuously take up lipids, for example by micropinocytosis.^47–52^ Lipids are a very heterogeneous group of macromolecules and parameters such as chain lengths and saturation of the acyl chains can have vastly different cellular effects including modulating inflammation and tumor immunity.^17,18^ We do not know whether BC cells deal with all lipids the same way. In this study, we observe comparable results using three distinct diets, 60% HFD, WD (45% kcal from fat), and KD (90% kcal from fat), which differ in their total fat content but have roughly similar fatty acid composition.

### KD and cancer

The idea of using dietary interventions as adjuvant therapy for cancer is now over a century old.^53^ Weight loss upon treatment completion is advised in women with obesity and a diagnosis of BC, since excessive weight is associated with a shorter time to recurrence and increased mortality.^3,4,32^ Yet not all weight loss interventions are equivalent in restoring metabolic health. Intermittent fasting and KD regimens have gained popularity as adjuvant cancer therapy.^54–56^ For example, while the ketogenic diet is a popular and effective diet to promote weight loss, there are concerns regarding the impact of this diet on lipid levels and cardiovascular health.^57^ However, KD are highly effective at suppressing glucose.^58^ Preclinical studies have shown positive results when tested in combination with targeted kinase inhibitors in *PIK3CA* mutant BC tumors, but clinical evidence for the efficacy of KDs is mixed and may be influenced by cancer subtype, genetic background, or a tumor-associated syndrome.^41,55^ We propose that individuals’ metabolic health and their tumor genotype should be considered when evaluating the efficacy of KDs, since cancer cells in patients with obesity may be reprogrammed to preferentially take up dietary lipids directly to fuel tumor growth, and KD, in this context, would provide additional lipid fuel. More broadly, we put forth that health nutrition plans as adjuvant therapies for cancer need to account for the lipid dependencies of cancer cells in the context of an individual’s metabolic health.

### Limitations

C57BL/6 mice are the standard model of diet-induced obesity, because of their propensity to rapidly gain weight upon HFD feeding and subsequently develop metabolic disorders. In contrast, immunocompromised nude mice, which are used for the study of human BC cell lines in an *in vivo* context, are resistant to diet-induced obesity.^59^ Accordingly, few validated models to study obesity-accelerated BC exist and our results are limited by the availability of relevant mouse BC cell lines. For example, no hormone positive C57BL/6 cell lines have been described to be accelerated by obesity so far and it remains to be determined whether our results also apply to hormone positive BC subtypes. We demonstrated that lowering systemic lipids via either diet-induced WL or pharmacological agent could restrict tumor growth, but whether more directly applicable mechanisms for human health (e.g. statins or GLP1 agonists) would have the same effect remains unknown. While our data show increased uptake of lipids in hyperlipidemia-accelerated BC, we do not yet know if only a subset of lipids drive tumorigenesis. Future investigation will establish if some fatty acids are less detrimental than others in promoting BC growth. Finally, an improved understanding of the mechanisms governing increased lipid acquisition and lipid fates within the tumor may identify targetable metabolic vulnerabilities in hyperlipidemia-accelerated BC.

## Materials and Methods

### Mice

This study used female wild type mice (**WT**) C57BL/6J (JAX #000664), B6.129P2-Apoe^tm1Unc/J^ (ApoE KO, JAX #002052) and B6.129S7-Ldlr^tm1Her/J^ (LDLR KO, JAX #002207). Upon reception at the University of Utah, mice were allowed to acclimatize for 2-3 weeks on a standard low-fat chow diet. Mice were 6-10 weeks old at the start of experimental dietary interventions. Mice were housed under a 12:12 light:dark cycle, with lights on at 6:00 am and had *ad libitum* access to food and water. All animal experiments were carried out in accordance with the guidelines and approved by the IACUC of the University of Utah. triolein

### Diets

All diets used in this study were obtained from Research Diets. Low fat diet (LFD, D12450K): 10% kcals from fat, 70% kcals from carbohydrates, 20% kcals from protein. Lard based 60% high fat diet (HFD, D12492): 60% kcals from fat, 20% kcals carb, 20% kcals protein. Lard based 90% fat ketogenic diet (KD, D16062902): 89.9% kcals from fat, 0.1% kcals from carbohydrates, 10% kcals from protein. Low fat medium protein control of the KD diet (KD-ctrl, D21110806): 10% kcals from fat, 80% kcals from carbohydrates, 10% kcals from protein. Western diet (WD, D12108C): 40% kcals from fat, 40% kcals from carbohydrates, 20% kcals from protein supplemented with 1.25% of cholesterol.

**Body composition** was measured using a Bruker Minispec small animal NMR (Bruker, Cat#LF50) in awake mice.

### Blood collection and analysis

Blood was collected by cheek bleeds either in the fasted state (5-6 h during the light phase), refed state (3-4 h after adding back food to fasted mice), or randomly fed state (before light onset) and indicated in the manuscript. Blood was collected in EDTA-coated tubes (Sarsdtet, 16.444.100) and serum separated by a 10-minute, 1000g spin. Triglycerides (TG), cholesterol (Cho) and non-esterified fatty acids (NEFA) were measured using colorimetric assays (Sekisui Diagnostics, TG: Cat #236-60, Chol: Cat #234-60, NEFA: Cat #:999-34691, 991-34891, 995-34791, 993-35191, 997-76491). Insulin levels were measured by Elisa (Crystal Chem, Cat #90080). Glucose was measured from the tail via a glucometer (Nova Max Plus).

### Cells and orthotopic injections

The E0771 breast tumor cell line of C57BL/6 origin (ATCC Cat #CRL-3461) was maintained in DMEM high glucose (Gibco Cat #11995040) supplemented with 10% FBS (Gibco Cat #A5670801), GlutaMax^TM^ (Gibco Cat#35050061) and 1% penicillin-streptomycin (Gibco Cat #15140122) at 37°C in a humidified 5% CO2 incubator. Cells were split two days before injection. E0771 (0.5 x 10^6^) were injected in 30µL Matrigel (Corning Cat #356231) bilaterally into the 4th mammary fat pads.

### Tumor measurements

Tumors were measured twice weekly using digital calipers (Fisherbrand Cat #S90187A) starting on day 7 post-tumor injection. Tumor volume was calculated using the following formula, (1/2 x length x width^2). At euthanasia, the tumors were dissected, cleaned and weighed on an analytical scale.

### ASOs preparation and injection

Antisense oligonucleotides (ASOs) targeting angiopoietin-like protein 3 (Angptl3)(Ionis #731875) and scramble control (Ionis #716837) were resuspended in PBS at a stock concentration of 35 mg/mL confirmed by absorbance spectrometry. A working solution at 2.5 mg/mL in PBS was used for subcutaneous injections at 25 mg/kg of body weight. Mice were treated with the ASOs on day 1, 3 and 7 of the first week of the intervention and then weekly. ^30^

### *In vivo* tumor lipid uptake assay

A lipid emulsion containing 1.25 µCi of ^3^H-triolein ([9,10-3H(N)]-triolein, Revvity) for every 150 µL of 5% Intralipid (Sigma-Aldrich, #68890-65-3) was prepared by mixing and tip sonication. ^60,61^ Mice bearing roughly 200-400 mg non necrotic tumors received a 150 µL oral bolus of the emulsion. Approximately 16 hours later, mice were euthanized and tumors and liver collected and weighed. Lipid extraction was performed using methyl tert-butyl ether (MTBE) and methanol. The organic and aqueous layers were transferred into separate scintillation vials containing Bio-Safe II scintillation cocktail (Research Products International, #111195). Samples were mixed thoroughly and counted using a Beckman Coulter LS6500 multi-purpose scintillation counter. The ^3^H counts in both fractions were added and the total counts normalized to the weight of the tissue to calculate ^3^H-triolein uptake per milligram of tissue. All experiments were carried out under protocols approved by the Radiation Safety Office of the University of Utah.

### Flow cytometry

E0771 cells were incubated with 100 nM C16-Bodipy (BODIPY™ FL C16, 4,4-Difluoro-5,7-Dimethyl-4-Bora-3a,4a-Diaza-s-Indacene-3-Hexadecanoic Acid, ThermoFisher, D3821) or 5 µg of pHrodo™ Red-LDL Conjugates (Low Density Lipoprotein from Human Plasma, pHrodo™ Red-LDL Conjugates, ThermoFisher, L34356) in normal growth media. After the indicated time, cells were thoroughly washed with PBS, collected and analyzed by flow cytometry using a BD FACSCanto™ Flow Cytometry System.

### Statistical Analysis

Results are expressed as mean with standard error to the mean (sem). Bar graphs show all individual points. A p-value of ≤ 0.05 was adopted as the cutoff for significance. Data normality was assessed using the Shapiro-Wilk test. A detailed description of the tests used for each figure panel is provided in the figure legend. GraphPad Prism was used for all statistical analyses.

## Funding

This work was supported by funds from NIDDK F32DK137475 to MRG, NIA R01AG065993 to AC, NCI UH2 CA286584 to AC, KIH and GSD, The American Cancer Society (DBG-23-1037804-01-TBE) to GSD, the V Foundation for Cancer Research to KIH, and the Pew Charitable Trusts to KIH.

## Acknowledgements

Some data reported in this publication were generated with assistance from the Flow Cytometry Core and the Metabolic Phenotyping Core of the University of Utah. We thank Elijah T. Matsuzaki, Seth Roberts and Ivan Delgado from the Chaix lab for assistance with animal feeding and tumor measurements and Annabel Lee for pilot cell culture experiments. We are grateful for seed funding from the Diabetes and Metabolism Research Center of the University of Utah and acknowledge the direct financial support provided by the Huntsman Cancer Foundation, the Breast and Gynecologic Cancers Disease Center, the Nuclear Control of Cell Growth and Differentiation and the Cell Response and Regulation Programs at the Huntsman Cancer Institute. We also acknowledge support by the National Cancer Institute of the National Institutes of Health under Award Number P30CA042014. The content is solely the responsibility of the authors and does not represent the official views of the NIH.

## Author contribution

Conceptualization: AC, KIH, GSD; Methodology: all authors; Data collection and analysis: all authors; Writing: all authors contributed to initial drat; final: AC, KIH, GSD; Funding and Resources: AC, KIH, GSD; Supervision: AC, KIH, GSD.

## Declaration of interests

The authors declare no competing interests.

## Notes

### Competing Interest Statement

The authors have declared no competing interest.

